# Isolation of Extracellular Vesicles from human plasma samples: The importance of controls

**DOI:** 10.1101/2022.10.31.514442

**Authors:** Migmar Tsamchoe, Stephanie Petrillo, Peter Metrakos, Anthoula Lazaris

**Affiliations:** Department of Anatomy and Cell Biology, McGill University, Quebec, Canada; Department of Surgery, McGill University Health Centre Research Institute, Quebec, Canada

**Keywords:** Extracellular vesicles, Isolation, Ultracentrifugation, Plasma, Thromboplastin

## Abstract

There are a number of methods for the isolation of extracellular vesicles (EV) which include the traditional ultracentrifugation to column-based kits available from different companies. Isolation of EVs from complex fluids, such as blood, has several challenges as the detection of low abundance molecules can easily be masked by more abundant proteins, when performing mass spectrometry. For this reason, several commercially available kits contain Thromboplastin D (TP-D) to promote clotting, thus removing clotting factors and abundant proteins resulting in increased detection of proteins. Our study demonstrates that plasma pretreated with Rabbit brain derived TP-D (the most common additive), generated a dynamic range of proteins compared to plasma alone, however, most of these proteins were contaminants introduced from the TP-D (99.1% purity). As an alternative, we tested recombinant TP and demonstrated that although it did not introduce any significant contaminants, we did not see any difference in the detection of proteins. Thus TP-D is not required, and any protein additives must be carefully screened.

## Introduction

Cells secrete membrane-enclosed vesicles (i.e., exosomes, microvesicles, apoptotic bodies) collectively known as extracellular vesicles (EV). EVs are found in a wide range of biological fluids, including blood, urine, saliva, amniotic fluid, and pleural fluid (1–6). There are two main groups of extracellular vesicles: exosomes of endosomal origin (40–100 nm in diameter) and shed vesicles (or ectosomes) pinched off from the plasma membrane (50–1000 nm in diameter). We refer to the collective group as EVs(7). EVs carry different biologically active molecules which will help in modulating the function of the targeted cells, thereby, facilitating the cell-cell communication. EVs are enriched with proteins, mRNAs, miRNA, lipids, metabolites, etc. providing potential genetic information and could be used for clinical applications (8).

There are a number of methodologies for isolating EVs from plasma (9), the classical method, allowing for the isolation of all size vesicles, is ultracentrifugation (10). Plasma is uniquely challenging as it is composed of many proteins, especially immunoglobulins and clotting factors which perturb the EV isolation leading to low recoveries of total EV number. For protein detection a common method is mass spectrometry, which is based on the ionization of molecules and can detect low abundance proteins in samples that are not complex. A challenge remains in increasing the ability to detect, identify, and quantify low abundant proteins. The presence of higher abundant species such as immunoglobulins, albumins, and non-EV lipid particles (such as lipoproteins) can mask the detection of low abundance proteins. Thus, a number of published protocols include thromboplastin (TP-D) to increase the clotting of the blood with the goal of eliminating the abundant proteins (11). There are commercially available methods for EV isolation such as miRCURY exosome kit, ExoQuick LP exosome isolation kits pre-treat the plasma with thrombin or thromboplastin. The purpose of this study was to assess the effect of different processing methods on EV isolation from plasma, specifically: the pre-treatment of plasma with thromboplastin from the most common source: rabbit brain extracts. EVs were isolated from 5 patients where we assessed 1. Plasma with TP-D used two sources: rabbit brain extract (rTP) vs human recombinant (huTP) 2. plasma alone. We also included additional groups where we followed the same protocol for EV isolation using 3. only PBS 4. PBS + TP-D, using the same two sources. Characterisations of the EVs were performed based on the MISEV guidelines (12). Analysis of the protein content was performed using mass spectrometry.

## Materials and Methods

### Human sample collection and preparation

Informed consent was obtained from all patients through the MUHC Liver Disease Biobank (LDB: MUHC research ethics board approved protocol REB#11-066-SDR). Bloods were collected in EDTA tubes, inverted immediately upon collection, and then spun at 2000 g at room temperature, for 15min.

### EV isolation using ultracentrifugation with and without thromboplastin

EVs were isolated using differential centrifugation with two ultra centrifugations. Briefly, we procured 20ml of plasma from 5 patients and split each patient sample into 4 tubes (5 mL/tube) and processed immediately. One tube was treated with rabbit brain tissue derived thromboplastin (Thromboplastin-D: HemosIL cat. 292273, with <0.9% rabbit brain tissue contaminant) and another tube with human recombinant Thromboplastin (Thermo fisher scientific). Next to remove cells, debris, Thromboplastins and Fibrins the plasma was centrifuge at 4700Xg for 15 mins followed by high-speed centrifugation (Beckman Avanti J-26XP) at 12,000 x g for 25mins and filtered using a 0.22μm filter (Ultident Scientific; Cat#229747). This was then followed by two sequential ultracentrifugation’s (Beckman Coulter) at 100,000 x g for 70mins. The pellet was resuspended with 0.1μm filtered DPBS (Multicell, cat 311-425-CL) between each spin. The final EV suspension was aliquoted and stored at -80C for further use. The same method was used however the plasma was replaced with the same volume of PBS alone, PBS with rabbit brain TP-D and PBS with human recombinant TP-D.

### Nanoparticle Tracking Analysis (NTA)

NTA was performed using a Malvern NanoSight NS500 instrument and NTA 3.1 Software. EVs were diluted in 0.1μm filtered DPBS and for each sample 3 videos were recorded with camera level 14-15 and detection threshold 5.

### Protein quantification

10μg of EV suspension was lysed with 10x RIPA (Sigma, Cat 20-188), and quantitated using Thermo Fisher Scientific micro BCA™ Protein Assay kit (catalog no. 23235) according to the manufacturer’s instruction using the Tecan microplate reader (Tecan Infinite 200 Pro).

### Western Blot Analysis

Equal concentration of proteins from EV samples (10μg) were loaded onto precast SDS-polyacrylamide gels (BioRad, Mini Protean TGX Cat# 456-1086) and transferred to Immobilon-E PVDF Transfer membrane (Sigma, IEVH85R). Blocked the membrane with 5% skimmed milk and incubated with primary antibody i.e., anti-TSG101 (Thermo Fisher Scientific, Cat# MA5-37764) and anti-TAPAI (Cat# ab79559) overnight at 4°C. After extensive washing of the membrane with 0.1% PBST, incubated with secondary antibody i.e., Goat anti-mouse IgG HRP conjugate (BioRad, cat# 170-6516). The blot is then added with ECL Prime western blotting detection reagent (Amersham™). For detection of the chemiluminescence with ImageQuant LAS4000 imager (GE life science).

### Proteomic analysis

Exosome proteins were loaded onto a single stacking gel band to remove detergents and salts. The gel band was reduced with DTT, alkylated with iodoacetic acid, and digested with trypsin. Extracted peptides were re-solubilized in 0.1% aqueous formic acid and loaded onto a Thermo Acclaim Pepmap (Thermo, 75uM ID X 2cm C18 3uM beads) precolumn and then onto an Acclaim Pepmap Easyspray (Thermo, 75uM X 15cm with 2μM C18 beads) analytical column separation using a Dionex Ultimate 3000 uHPLC at 220 nl/min with a gradient of 2-35% organic (0.1% formic acid in acetonitrile) over 3 hours. Peptides were analyzed using a Thermo Orbitrap Fusion mass spectrometer operating at 120,000 resolutions (FWHM in MS1) with HCD sequencing at top speed (15,000 FWHM) of all peptides with a charge of 2+ or greater. The raw data were converted into *.mgf format (Mascot generic format), searched using the GPM X! Tandem against Swissprot Human 2018 protein FASTA files. The database search results were loaded onto Scaffold Q+ Scaffold_4.4.8 (Proteome Sciences) for statistical treatment and data visualization.

FunRich: The protein list from the scaffold were filtered at Protein threshold of 95% and peptide ≥2. The list is then loaded in the FunRich analysis tool which have EV database called Vesiclepedia incorporated within the tool and number of EV derived proteins were visualised using Venn diagram.

### Transmission Electron Microscopy

For gold labelling TEM analysis, EVs are washed with 0.1% sodium cacodylate and fixed with 2.5% glutaraldehyde fixative solution. Next, we treated the EV sample with BCO for 5 mins followed by incubation in the primary antibody overnight at 4°C. The sample was washed twice with distilled water and then incubated in secondary antibody for 30mins. After two washes with distilled water the sample was stained with 4% uranyl acetate for 3mins. The dried grid was then examined under Tecnai 12 BioTwin 120kV TEM at the Facility for Electron Microscopy Research (FEMR).

### miRNA Isolation and detection

EVs were first treated with phenol-chloroform, followed by miRNA isolation using the total exosome protein and RNA isolation kit (Thermoscientific). All the steps were followed according to the manufacturer’s instruction. The miRNA concentration and quality were checked using Agilent 6000 pico-kit 5067-1513 and analysed using Bioanalyzer software.

### Statistical Analysis

Used GraphPad Prism 9.4.1 software for plotting the graph. Prism is used for performing mean and standard deviation calculation.

## Results

### Comparison between plasma derived EVs isolation using 2UC with and without Rabbit brain derived thromboplastin (rTP)

We used the two ultracentrifugation method adapted from Choi et.al (Fig 1) (13) to isolate EVs. To study whether addition of rTP would help in enriching EVs from plasma sample, five plasma samples was treated with rTP and compared to a second group of 5 plasma samples without rTP. For the experimental control, instead of plasma we used PBS, treated with rTP and PBS without rTP, and processed the EV isolation simultaneously. As reported by others we observed an increase in the total number of EV particles in the plasma with rTP sample group compared to the plasma alone (Fig 2a) and observed ∼4.3fold increase in the protein yield. Plasma alone group had an average concentration of 44.5ug/ml versus average of 195ug/ml in plasma with rTP. Surprisingly, an average of 1.60E+11 particles (Fig 2b) was detected in the control sample (PBS + rTP alone sample) with a protein concentration of 16.9ug/ml. The PBS alone sample had no detectable EVs. The increase in protein concentration was validated by western blot using TSG101 and CD81. All samples showed the presence of both markers (Fig 2c). Electron microscope was used to visualise the EV in the isolated sample (Fig2d). To further investigate the type and number of proteins detected, we performed LCMS/MS on all samples. We detected 231 proteins in the plasma alone, 826 proteins in the plasma+rTP and surprisingly, 864 proteins in the PBS+rTP. Using FunRich analysis we cross referenced the EV data bank (i.e., Vesiclepedia) with that of our dataset. We identified 183 proteins in plasma alone, 727 proteins in plasma+rTP and 752 proteins in PBS+rTP EVs overlapping with Vesiclepedia (Fig 2e). As human and rabbit proteins have high similarities it is very difficult to identify the contaminating rabbit proteins and exclude them from the analysis. The scaffold data indicates clearly that the rTP is introducing many contaminating proteins.

**Fig1a:**
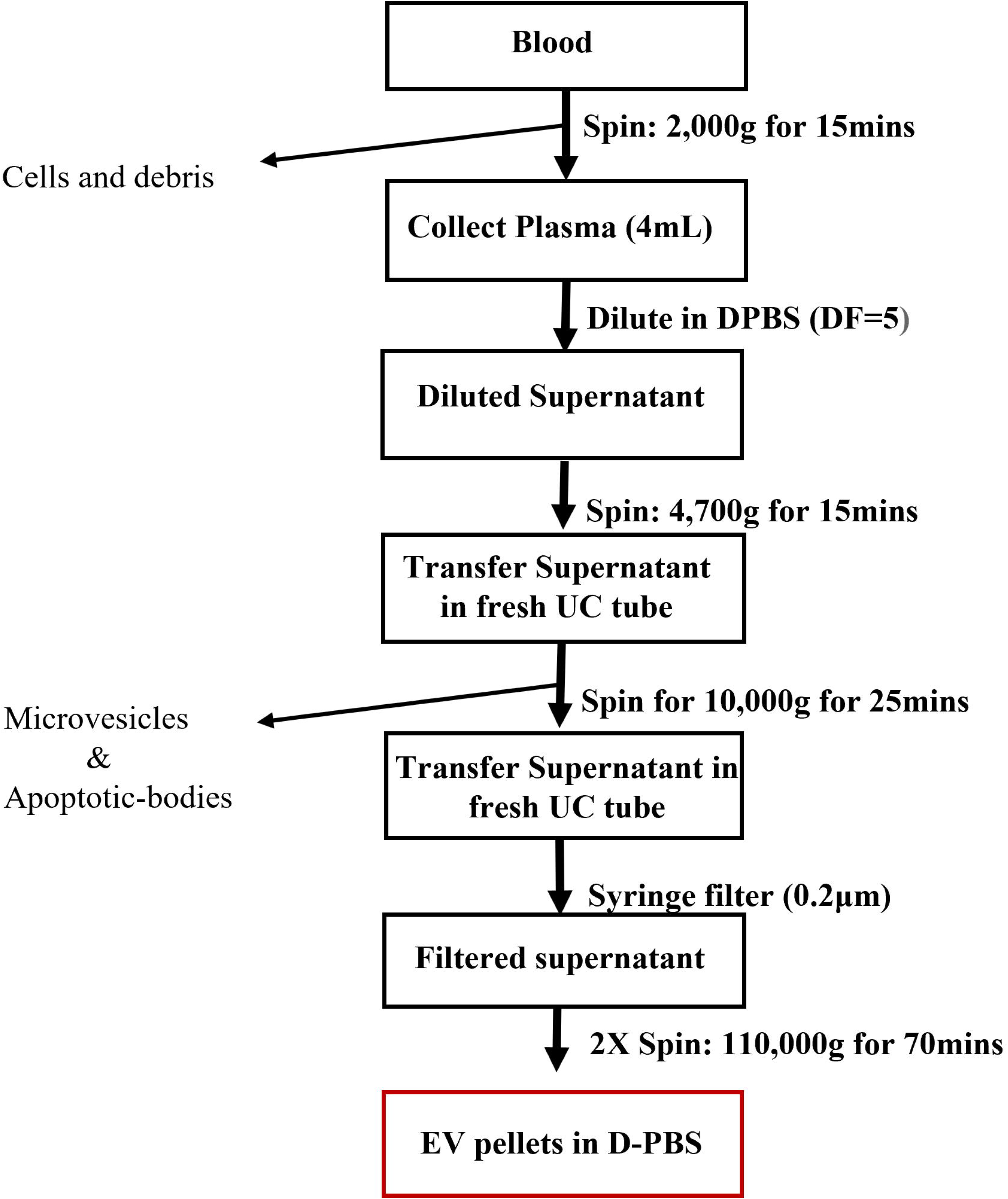
Two Ultracentrifugation method of Extracellular vesicle Isolation from plasma.

A similar trend was observed for EV derived miRNAs. The bio-analyzer results demonstrated miRNA detection in all the samples i.e., the plasma alone, plasma +rTP and PBS+rTP. Thus, resulting in contributing miRNA contaminants from the rTP during EV isolation from the plasma sample (Fig 2f). It is also important to note that, like rabbit proteins, miRNAs are highly conserved throughout species, therefore, it is impossible to differentiate the origin of the miRNA.

**Fig2.**
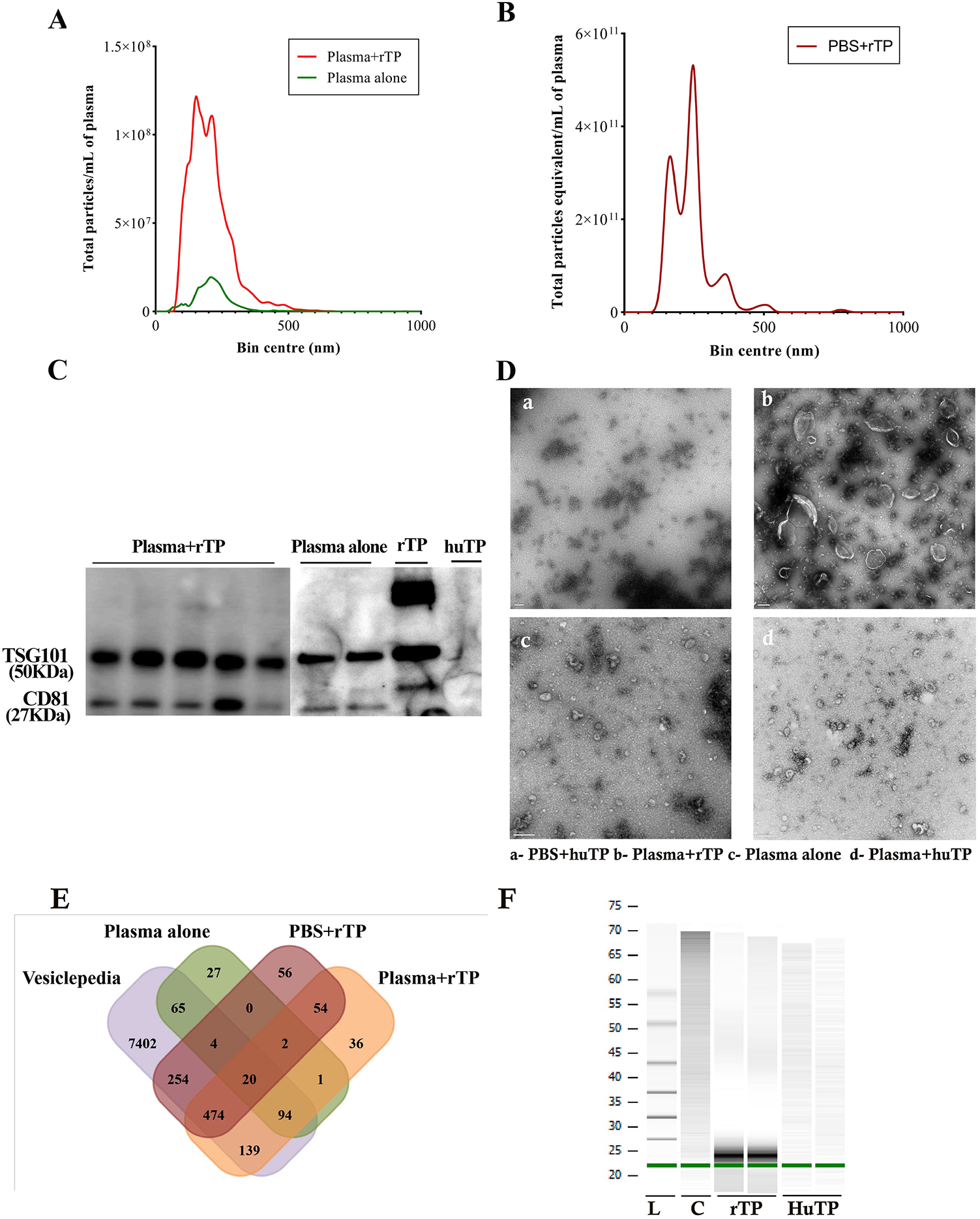
Comparative study of Ultracentrifugation EV isolation (Plasma alone) with Ultracentrifugation method treated with rabbit brain derived Thromboplastin (Plasma+rTP): **a**. NTA curve of Plasma alone vs Plasma+rTP and **b**. PBS+rTP showing total particles/mL of plasma against particle size in nm. Curves represent the average of 5 samples. **c**. Western blot showing EV marker in Plasma+rTP, Plasma alone and PBS+ rTP. **d**. Electron microscope image showing presence of EV. **e**. Venn diagram showing total shared EV proteins of Vesiclepedia with that of Plasma alone, Plasma+rTP and PBS+rTP. **f**. Bioanalyser result showing miRNA contaminants from plasma+rTP.

### Human Recombinant Thromboplastin for the EV isolation from plasma

Having identified that the rabbit brain derived TP is introducing significant contaminants; we decided to assess a pure form of TP, human recombinant TP (huTP), with the goal of promoting clotting and eliminating clotting factors and abundant proteins, eventually allowing the detection of less abundant proteins by mass spectrometry. The same 3 groups were evaluated: plasma alone, plasma + huTP and PBS + huTP. We observed no significant difference in the number of particles between the plasma and plasma+ huTP (Fig 3a). Furthermore, the PBS+huTP showed below optimum detectable particle numbers (Fig 3a). We then analyzed the total number of proteins from LCMS/MS using scaffold and observed a similar number of total proteins detected (i.e., plasma alone 183 and plasma+huTP 189 proteins). Most of the proteins between the huTP treated and non-treated plasma samples were similar based on the FunRich analysis, except for a small number of proteins with very low spectral counts that were mostly Keratins, known contaminants in LCMS/MS. FunRich analysis detected approximately the same number of EV proteins with no significant differences (Fig 3b). Even though PBS+huTP sample showed below the levels of detection for the BCA, we nevertheless processed the samples for LCMS/MS. Most of the proteins detected in the huTP alone were keratins, Immunoglobulins, and albumins, which are common contaminants. FunRich analysis yield 38 overlapping proteins with Vesiclepedia (Fig 3b) but most of these proteins had very low spectral count (Fig 3c).

**Fig3.**
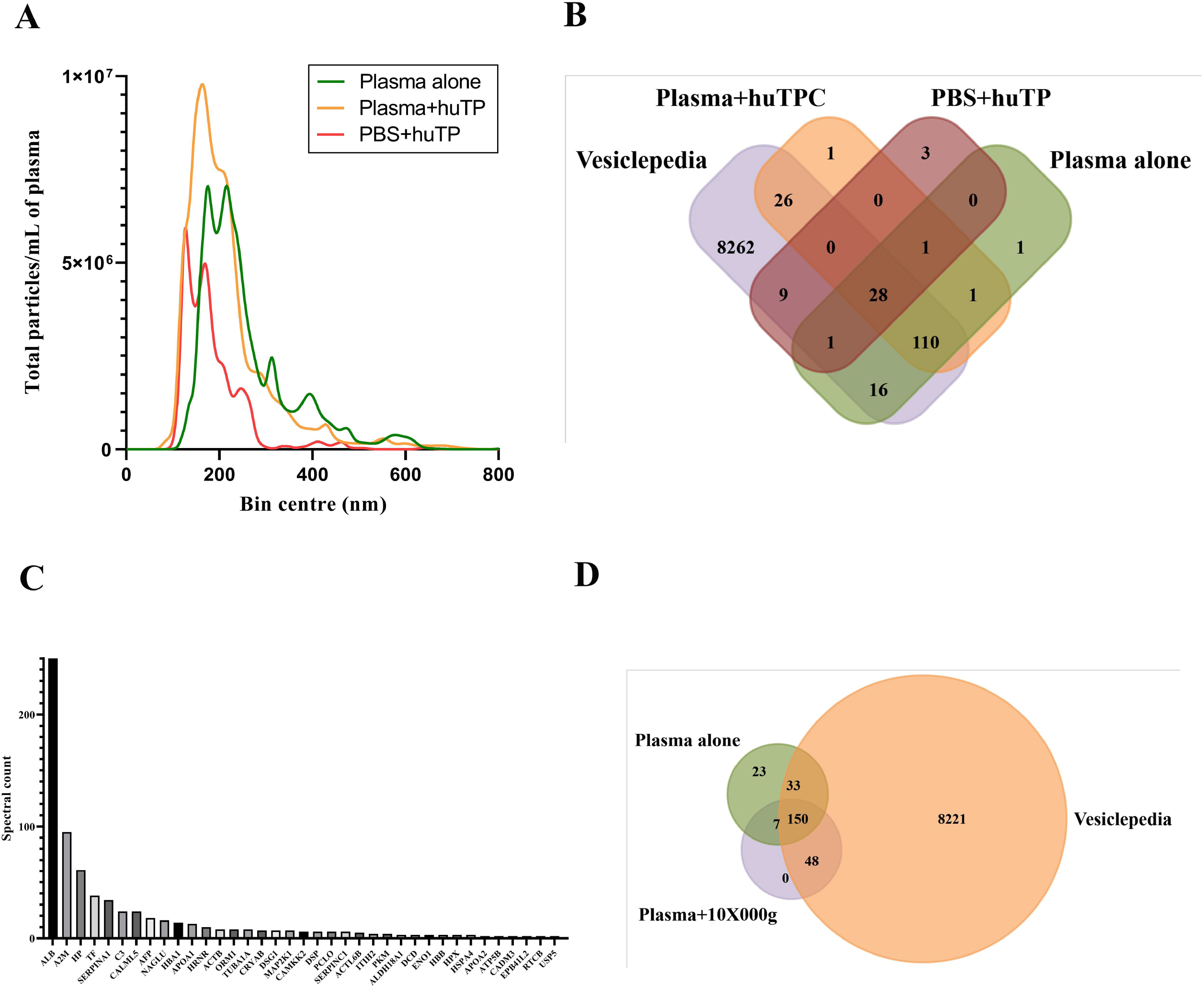
Comparative study of Ultracentrifugation method (Plasma alone) vs plasma with human recombinant thromboplastin (Plasma+huTP): **a**. NTA curve of Plasma alone, Plasma+huTP,PBS+ huTP showing Total particles/mL plasma against particle size in nm **b**. Venn diagram showing shared EV in Vesiclepedia database with proteins detected in Plasma alone, PBS+huTP and Plasma+huTP **c**. PBS+huTP graph showing list of proteins detected vs spectral count **d**. Venn diagram showing shared EV in Vesiclepedia database with proteins detected in plasma with and without extra 10×000g spin. Abbreviations- (L=Ladder and C=Control)

Furthermore, several publications perform an extra spin of 10,000g on the plasma sample, a procedure that is not commonly performed on plasma samples from biobanks. We therefore compared EV isolations with and without the additional spin and compared freshly processed samples to frozen samples. Mass spectrometry for five patient samples in each group of plasma with and without the extra 10,000g spin detected more proteins (i.e., 213 proteins) in the sample without the extra spin. Further, plasma alone have 85% and plasma with 10,000g have 96% proteins common with the Vesiclepedia data base with 10% of unique proteins observed in plasma alone while no unique proteins detected in plasma with extra spin. (Fig3d).

## Discussion

The focus on finding non-invasive potential disease biomarkers using circulating vesicles had gained popularity in the past few decades (14–18). Although there has been a plethora of publications on EVs, it is important to choose a method of isolation that is reproducible and does not introduce contaminants. In this study, we wished to assess the traditional ultracentrifugation method and additives that are commonly used and included in existing commercial kits. We were very surprised by our findings that the addition of rabbit brain derived thromboplastin, used in several commercial kits and publications (2,7–9,12–14,19–50) introduce EV contaminants which can potentially result in misleading findings, wasting resources and time. As an alternative, we used human recombinant TP, with the goal of having a pure source with few or no animal contaminants to test if the addition of TP is beneficial to increasing the recovery of EVs. Interestingly as there was no change in the EV yield and EV proteome profile, we concluded that the huTP does not introduce contaminants and does not improve in the detection of proteins and thus the addition of TP is unnecessary.

As liquid biopsy, specifically plasma, are becoming common fluids to screen patients, the collection and processing of the blood sample from multi-sites may introduce variation between the samples. We therefore decided to evaluate if fresh vs frozen samples will generate different protein or miRNA profiles. Our data clearly demonstrated no significant differences in protein content whether the plasma samples were processed fresh or frozen. In addition, as multiple labs include an extra processing step that is not feasible to introduce into standard biobanking processes, we also evaluated the effect of the extra spin (10,000g) on the protein and miRNA profiles. Once again, we observed no significant difference indicating that bio-banked plasma specimens can be used reproducibly across multiple sites.

Our study highlights the need to include a proper control in all EV isolation methods, as any traces of animal sample contaminants such as <0.9% of rabbit brain derived tissue TP could have dramatic effect on the results.

## Acknowledgement

The authors like to acknowledge Dr. Janusz Rak and Dr. Tommy Nilsson for their constant guidance, Lotfi Amri, Jaclyn Chabot, and Kurt Dejgaard for their support. The project was supported by MEDTEQ innovation for Health for the project titled “AI-powered multi-omic signature discovery for precision care of patients with Colorectal Cancer Liver Metastases (CRCLM)” under the grant number MEDTEQ 11-N CRCLM Biomarkers. We would like to also acknowledge the MUHC Liver Disease Biobank and thank all the patients who participated in this study and providing their precious blood samples which played a key role in successfully completing this research.

## Author Contributions

M.T., P.M., and A.L., designed the study and wrote the manuscript. M.T., and L.A performed the experiments, S.P. collected the patient samples and consents, M.T. analysed the data, generated the images, and interpreted the results, A.L. and P.M provided feedback and provided guidance throughout the study.

## Supporting Information

**S1 Excel : List of Proteins detected with spectral count information**.

## References

1. Bard MP, Hegmans JP, Hemmes A, Luider TM, Willemsen R, Severijnen L-AA, et al. Proteomic analysis of exosomes isolated from human malignant pleural effusions. Am J Respir Cell Mol Biol. 2004;31(1):114–21.

2. Runz S, Keller S, Rupp C, Stoeck A, Issa Y, Koensgen D, et al. Malignant ascites-derived exosomes of ovarian carcinoma patients contain CD24 and EpCAM. Gynecol Oncol. 2007;107(3):563–71.

3. Michael A, Bajracharya SD. Yuen PST, Zhou H, Star RA. Illei GG. et al. Exosomes from human saliva as a source of microRNA biomarkers. Oral Dis. 2010;16(1):34–8.

4. Palanisamy V, Sharma S, Deshpande A, Zhou H, Gimzewski J, Wong DT. Nanostructural and transcriptomic analyses of human saliva derived exosomes. PLoS One. 2010;5(1):e8577.

5. Keller S, Ridinger J, Rupp A-K, Janssen JWG, Altevogt P. Body fluid derived exosomes as a novel template for clinical diagnostics. J Transl Med. 2011;9(1):1–9.

6. Street JM. Barran PE. Mackay CL. Weidt S, Balmforth C, Walsh TS. et al. Identification and proteomic profiling of exosomes in human cerebrospinal fluid. J Transl Med. 2012;10(1):1–7.

7. Simpson RJ. Kalra H, Mathivanan S. ExoCarta as a resource for exosomal research. J Extracell vesicles. 2012;1(1):18374.

8. Crawford N. The Presence of Contractile Proteins in Platelet Microparticles Isolated from Human and Animal Platelet□free Plasma. Br J Haematol. 1971;21(1):53–69.

9. Willms E, Cabañas C, Mäger I, Wood MJA, Vader P. Extracellular vesicle heterogeneity: subpopulations, isolation techniques, and diverse functions in cancer progression. Front Immunol. 2018;9:738.

10. Crigna AT. Fricke F, Nitschke K, Worst T, Erb U, Karremann M, et al. Inter-Laboratory Comparison of Extracellular Vesicle Isolation Based on Ultracentrifugation. Transfus Med Hemotherapy. 2021;48(1):48–59.

11. Quackenbush JF. Cassidy PB. Pfeffer LM. Boucher KM. Hawkes JE. Pfeffer SR. et al. Isolation of circulating microRNAs from microvesicles found in human plasma. Methods Mol Biol. 2014;1102:641–53.

12. Théry C, Witwer KW. Aikawa E, Alcaraz MJ. Anderson JD. Andriantsitohaina R, et al. Minimal information for studies of extracellular vesicles 2018 ( MISEV2018 ): a position statement of the International Society for Extracellular Vesicles and update of the MISEV2014 guidelines. 2018;3078.

13. Choi D-S, Gho YS. Isolation of extracellular vesicles for proteomic profiling. In: Proteomic Profiling. Springer; 2015. p. 167–77.

14. De Toro J, Herschlik L, Waldner C, Mongini C. Emerging roles of exosomes in normal and pathological conditions: new insights for diagnosis and therapeutic applications. Front Immunol. 2015;6:203.

15. Kim IA. Hur JY. Kim HJ. Lee SE. Kim WS. Lee KY. Liquid biopsy using extracellular vesicle–derived DNA in lung adenocarcinoma. J Pathol Transl Med. 2020;54(6):453.

16. de Miguel Pérez D, Martínez AR. Palomo AO. Ureña MD. Puche JLG, Remacho AR. et al. Extracellular vesicle-miRNAs as liquid biopsy biomarkers for disease identification and prognosis in metastatic colorectal cancer patients. Sci Rep. 2020;10(1):1–13.

17. Zhou B, Xu K, Zheng X, Chen T, Wang J, Song Y, et al. Application of exosomes as liquid biopsy in clinical diagnosis. Signal Transduct Target Ther. 2020;5(1):1–14.

18. Desmond BJ. Dennett ER. Danielson KM. Circulating Extracellular Vesicle MicroRNA as Diagnostic Biomarkers in Early Colorectal Cancer—A Review. Cancers (Basel). 2020;12(1):52.

19. Winston CN. Goetzl EJ. Akers JC. Carter BS. Rockenstein EM. Galasko D, et al. Prediction of conversion from mild cognitive impairment to dementia with neuronally derived blood exosome protein profile. Alzheimer’s Dement Diagnosis, Assess Dis Monit. 2016;3:63–72.

20. Davidson S. Microvesicles and Exosomes in Local and Distant Communication with the Heart. In: Stem Cells and Cardiac Regeneration. Springer; 2016. p. 143–62.

21. Rabinowits G, Bowden M, Flores LM. Verselis S, Vergara V, Jo VY. et al. Comparative analysis of microRNA expression among benign and malignant tongue tissue and plasma of patients with tongue cancer. Front Oncol. 2017;7:191.

22. Sun B, Dalvi P, Abadjian L, Tang N, Pulliam L. Blood neuron-derived exosomes as biomarkers of cognitive impairment in HIV. AIDS. 2017;31(14):F9.

23. Du M, Giridhar K V, Tian Y, Tschannen MR. Zhu J, Huang C-C, et al. Plasma exosomal miRNAs-based prognosis in metastatic kidney cancer. Oncotarget. 2017;8(38):63703.

24. Chae M-S, Kim J, Jeong D, Kim Y, Roh JH. Lee SM. et al. Enhancing surface functionality of reduced graphene oxide biosensors by oxygen plasma treatment for Alzheimer’s disease diagnosis. Biosens Bioelectron. 2017;92:610–7.

25. Eitan E, Green J, Bodogai M, Mode NA. Bæk R, Jørgensen MM. et al. Age-related changes in plasma extracellular vesicle characteristics and internalization by leukocytes. Sci Rep. 2017;7(1):1–14.

26. Goetzl EJ. Abner EL. Jicha GA. Kapogiannis D, Schwartz JB. Declining levels of functionally specialized synaptic proteins in plasma neuronal exosomes with progression of Alzheimer’s disease. FASEB J. 2018;32(2):888–93.

27. Patterson SA. Deep G, Brinkley TE. Detection of the receptor for advanced glycation endproducts in neuronally-derived exosomes in plasma. Biochem Biophys Res Commun. 2018;500(4):892–6.

28. Gill J, Mustapic M, Diaz-Arrastia R, Lange R, Gulyani S, Diehl T, et al. Higher exosomal tau, amyloid-beta 42 and IL-10 are associated with mild TBIs and chronic symptoms in military personnel. Brain Inj [Internet]. 2018;32(10):1277–84. Available from: https://doi.org/10.1080/02699052.2018.1471738

29. Goetzl L, Merabova N, Darbinian N, Martirosyan D, Poletto E, Fugarolas K, et al. Diagnostic potential of neural exosome cargo as biomarkers for acute brain injury. Ann Clin Transl Neurol. 2018;5(1):4–10.

30. Stranska R, Gysbrechts L, Wouters J, Vermeersch P, Bloch K, Dierickx D, et al. Comparison of membrane affinity-based method with size-exclusion chromatography for isolation of exosome-like vesicles from human plasma. J Transl Med. 2018;16(1):1–9.

31. Moon S, Shin DW. Kim S, Lee Y-S, Mankhong S, Yang SW. et al. Enrichment of exosome-like extracellular vesicles from plasma suitable for clinical vesicular miRNA biomarker research. J Clin Med. 2019;8(11):1995.

32. Goetzl EJ. Elahi FM. Mustapic M, Kapogiannis D, Pryhoda M, Gilmore A, et al. Altered levels of plasma neuron□derived exosomes and their cargo proteins characterize acute and chronic mild traumatic brain injury. FASEB J. 2019;33(4):5082–8.

33. Mustapic M, Tran J, Craft S, Kapogiannis D. Extracellular Vesicle Biomarkers Track Cognitive Changes Following Intranasal Insulin in Alzheimer’s Disease. J Alzheimer’s Dis. 2019;69(2):489–98.

34. Sun B, Fernandes N, Pulliam L. Profile of neuronal exosomes in HIV cognitive impairment exposes sex differences. Aids. 2019;33(11):1683–92.

35. Gu D, Liu F, Meng M, Zhang L, Gordon ML. Wang Y, et al. Elevated matrix metalloproteinase□9 levels in neuronal extracellular vesicles in Alzheimer’s disease. Ann Clin Transl Neurol. 2020;7(9):1681–91.

36. Liang M, Yu S, Tang S, Bai L, Cheng J, Gu Y, et al. A Panel of Plasma exosomal miRNAs as potential biomarkers for differential diagnosis of thyroid nodules. Front Genet. 2020;11:449.

37. Pulliam L, Liston M, Sun B, Narvid J. Using neuronal extracellular vesicles and machine learning to predict cognitive deficits in HIV. J Neurovirol. 2020;26(6):880–7.

38. Zhang N, Gu D, Meng M, Gordon ML. TDP-43 Is Elevated in Plasma Neuronal-Derived Exosomes of Patients With Alzheimer’s Disease. Front Aging Neurosci. 2020;12(June):1– 8.

39. Goetzl EJ. Peltz CB. Mustapic M, Kapogiannis D, Yaffe K. Neuron-derived plasma exosome proteins after remote traumatic brain injury. J Neurotrauma. 2020;37(2):382–8.

40. Goetzl EJ. Srihari VH. Guloksuz S, Ferrara M, Tek C, Heninger GR. Decreased mitochondrial electron transport proteins and increased complement mediators in plasma neural-derived exosomes of early psychosis. Transl Psychiatry [Internet]. 2020;10(1). Available from: http://dx.doi.org/10.1038/s41398-020-01046-3

41. Goetzl EJ. Yaffe K, Peltz CB. Ledreux A, Gorgens K, Davidson B, et al. Traumatic brain injury increases plasma astrocyte-derived exosome levels of neurotoxic complement proteins. FASEB J. 2020;34(2):3359–66.

42. Wei F, Wang A, Wang Q, Han W, Rong R, Wang L, et al. Wei et al., 2020. 2020;12(12):12002–18.

43. Hornung S, Dutta S, Bitan G. CNS-Derived Blood Exosomes as a Promising Source of Biomarkers: Opportunities and Challenges. Front Mol Neurosci. 2020;13(March):1–16.

44. Lazo S, Noren N, Jamal H, Erez G, Mode NA. Alan QL. et al. Mitochondrial DN. in extracellular vesicles declines with age. 2021;(April 2020):1–15.

45. de Oliveira Jr GP. Porto WF. Palu CC. Pereira LM. Reis AMM, Marçola TG. et al. Effects of endurance racing on horse plasma extracellular particle miRNA. Equine Vet J. 2021;53(3):618–27.

46. Goetzl EJ. Srihari VH. Guloksuz S, Ferrara M, Tek C, Heninger GR. Neural cell□derived plasma exosome protein abnormalities implicate mitochondrial impairment in first episodes of psychosis. FASEB J. 2021;35(2):e21339.

47. Peluso MJ. Deeks SG. Mustapic M, Kapogiannis D, Henrich TJ. Lu S, et al. SARS-CoV-2 and Mitochondrial Proteins in Neural-Derived Exosomes of COVID-19. 2022;1–10.

48. Michael A, Bajracharya SD. Yuen PST, Zhou H, Star RA. Illei GG. et al. Exosomes from human saliva as a source of microRNA biomarkers. Oral Dis. 2010;16(1):34–8.

49. Kanwar SS. Dunlay CJ. Simeone DM. Nagrath S. Microfluidic device (ExoChip) for on- chip isolation, quantification and characterization of circulating exosomes. Lab Chip. 2014;14(11):1891–900.

50. Goetzl EJ. Kapogiannis D, Schwartz JB. Lobach I V, Goetzl L, Abner EL. et al. Decreased synaptic proteins in neuronal exosomes of frontotemporal dementia and Alzheimer’s disease. FASEB J. 2016;30(12):4141–8.

